# Learning polygenic scores for human blood cell traits

**DOI:** 10.1101/2020.02.17.952788

**Authors:** Yu Xu, Dragana Vuckovic, Scott C Ritchie, Parsa Akbari, Tao Jiang, Jason Grealey, Adam S. Butterworth, Willem H Ouwehand, David J Roberts, Emanuele Di Angelantonio, John Danesh, Nicole Soranzo, Michael Inouye

## Abstract

Polygenic scores (PGSs) for blood cell traits can be constructed using summary statistics from genome-wide association studies. As the selection of variants and the modelling of their interactions in PGSs may be limited by univariate analysis, therefore, such a conventional method may yield sub-optional performance. This study evaluated the relative effectiveness of four machine learning and deep learning methods, as well as a univariate method, in the construction of PGSs for 26 blood cell traits, using data from UK Biobank (n=~400,000) and INTERVAL (n=~40,000). Our results showed that learning methods can improve PGSs construction for nearly every blood cell trait considered, with this superiority explained by the ability of machine learning methods to capture interactions among variants. This study also demonstrated that populations can be well stratified by the PGSs of these blood cell traits, even for traits that exhibit large differences between ages and sexes, suggesting potential for disease prevention. As our study found genetic correlations between the PGSs for blood cell traits and PGSs for several common human diseases (recapitulating well-known associations between the blood cell traits themselves and certain diseases), it suggests that blood cell traits may be indicators or/and mediators for a variety of common disorders via shared genetic variants and functional pathways.

## Introduction

Blood cells play essential roles in a variety of biological processes, such as oxygen transport, iron homeostasis, and pathogen clearance^1–3^. Abnormalities in blood cell traits, such as the number of cells, the proportions of different types, their sizes and morphology and so their likely functions, have been associated with a range of human diseases, such as reticulocyte indices with coronary heart disease^4^, or eosinophil counts with asthma^5^.

Blood cell traits are heritable, and their genetic architecture has been found to be highly polygenic. Analysis of the UK Biobank (UKB)^6,7^ and INTERVAL^8^ cohorts have suggested that between 18% to 30% of the variance in erythrocyte counts and morphology can be explained by hundreds of common autosomal variants^4^. It is expected, therefore, that levels of these traits can to some extent be predicted by genetic variants via polygenic scores (PGSs)^9^. Blood cell traits can be considered part of intermediary biological processes which may be involved in pathogenesis, and thus PGSs for blood cell traits may help better identify innate inter-individual differences in these traits, those which are related to disease risk, and those which may help identify novel therapeutic targets^10^.

A PGS is most commonly constructed as a weighted sum of genetic variants, typically single nucleotide polymorphisms (SNPs), carried by an individual, where the genetic variants are selected and their weights quantified via univariate analysis in a corresponding genome-wide association study (GWAS)^9,11^. As is the nature of univariate analysis, this conventional PGS method largely relies on hard cut-off thresholds to identify associated variants at the population level, e.g. *p*-value thresholding for selection of significant variants and *r*^*2*^ thresholding for selection of independent variants^12^. However, usage of harder or softer thresholds would either lead to weakly predictive PGSs, or bring in unrelated and/or over-representative variants, e.g. variants in high linkage disequilibrium (LD), that could weaken predictive power^13^. In practice, it is challenging to find an ideal set of thresholds that identify the associated variants from millions of other variants for maximal prediction power of a given trait or disease. Thus, PGSs obtained through this method are likely under-predictive.

A further limitation of the conventional PGS approach is its inherent assumption of linearity. Estimation of the weight of each variant alone through univariate association tests leaves no modelling considerations for joint effects between variants. However, studies have shown that variants, for example in the MHC region, can exhibit significant non-linear effects on traits/diseases through interactions^14–18^. Thus, in particular for traits of complex architecture, there may be considerable scope for improvement in genomic prediction compared to the linear univariate approach taken to construct most previous PGSs^18–20^.

Machine learning techniques have demonstrated superiority in constructing polygenic scores for several traits and diseases, largely immune-related, e.g. celiac disease, type 1 diabetes, and Crohn’s disease^19–23^. Owing to their simplicity and efficiency, multivariate linear models, such as elastic net and support vector machines, are one of the most widely used categories of machine learning methods for PGS construction. Although these approaches also entail assumptions of linearity, they may implicitly model interactions amongst variants via effect size shrinkage and variant selection techniques, e.g. L1 and L2 norms^20^. Non-linear models, e.g. polynomial regression, provide an explicit way to model the mutual interactions among input features; however, traditional non-linear learning methods usually suffer from poor efficiency and do not scale effectively when applied to high-dimensional genomic data. The rapid development of deep learning methods, e.g. Multilayer Perceptrons (MLPs) and Convolutional Neural Networks (CNNs), known for their power to model a wide spectrum of linear and non-linear correlations, and their supporting hardware architectures, e.g. Graphical Processing Units (GPUs) and Tensor Processing Units (TPUs), allow us to train complex non-linear models on large datasets within a reasonable timeframe and computing power.

This study aims to evaluate the relative effectiveness of the machine learning, deep learning and univariate methods in construction of PGSs for blood cell traits (see **Figure 1** for study workflow). We investigate three key questions: 1) To what extent can machine learning or deep learning methods improve construction of PGSs for blood cell traits? 2) What potential insights into the genetic architecture of blood cell traits and its relevance to disease can learned PGSs offer? and 3) Can PGSs be used to stratify trajectories of blood cell traits? To answer these questions, we examined the relative performances of the univariate PGS method, two linear machine learning methods that have different regularization techniques (elastic net and Bayesian ridge), and two deep learning methods that can model complex linear and non-linear correlations, namely Multilayer Perceptrons (MLPs) and Convolutional Neural Networks (CNNs). Taken together, we construct and externally validate PGSs for 26 blood cell traits, and make these PGSs available to the community via the PGS Catalog^24^.

**Figure 1.**
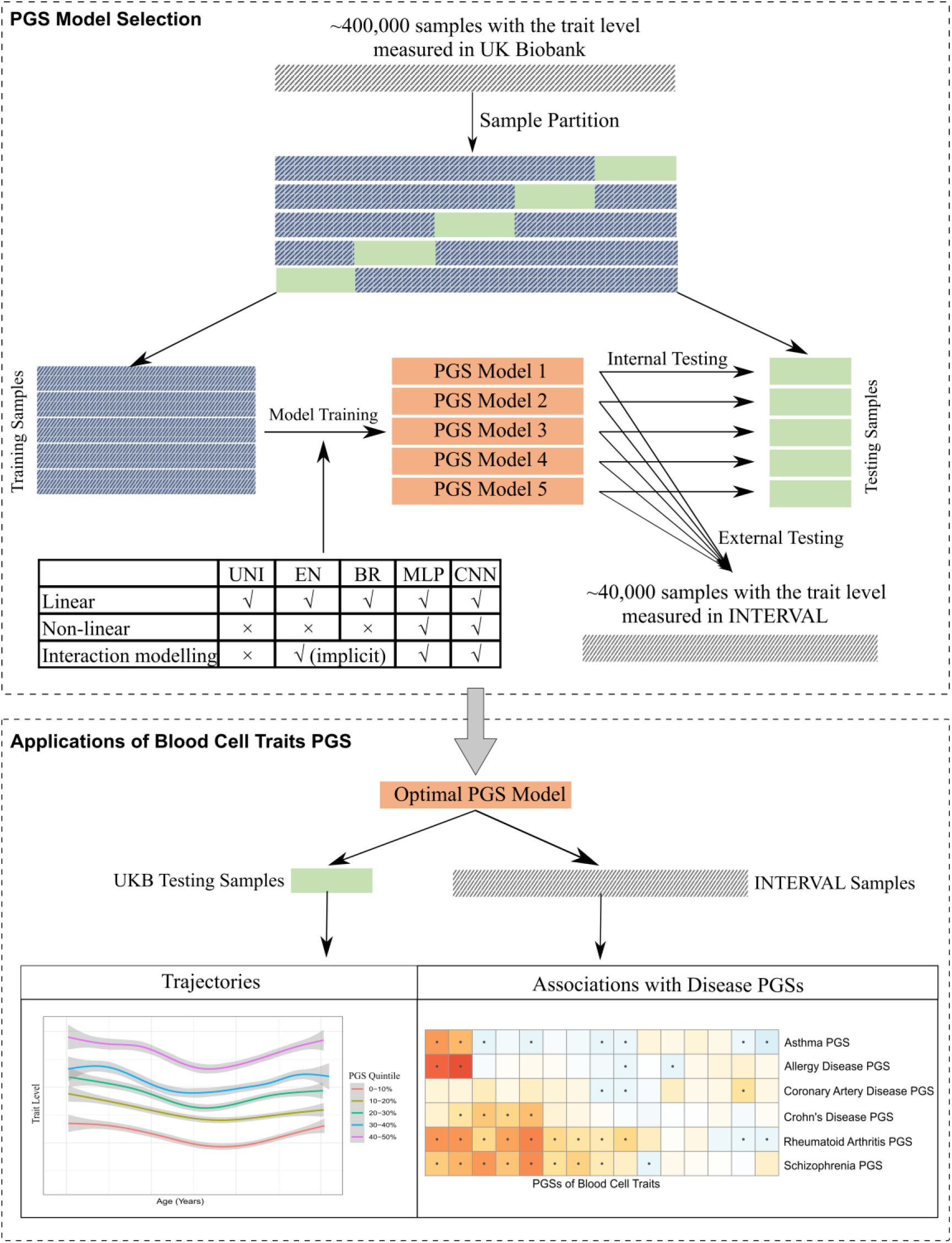
The study workflow of PGS construction of blood cell traits using learning methods and its applications. Five PGS methods were evaluated in this study: univariate method (UNI), elastic net (EN), Bayesian ridge (BR) and multilayer precepton (MLP) and convolutional neural network (CNN).

## Data and Methods

### Data and Quality Control

This study analysed 26 different traits across three blood cell types: platelets, red blood cells, and white blood cells (Table S1 and Table S2) that were measured in UK Biobank^6,7^ and INTERVAL^8^ cohorts. The UK Biobank is a cohort including 500,000 individuals living in the UK who were recruited between 2006 and 2010, aged between 40 and 69 years at recruitment. INTERVAL is a randomised trial of 50,000 healthy blood donors, aged 18 years or older at recruitment. As construction and evaluation of PGSs are highly dependent on the quality of both phenotype and genotype data used, we adopted the established protocols described in the previous work^4^, adjusting measured values for blood cell trait values to help account for a variety of environmental and technical factors, as well as the first 10 genetic principal components. Technical variables include the time between venepuncture and full blood cell analysis, seasonal effects, centre of sample collection, time dependent drift of equipment, systematic differences in equipment; environmental variables include sex, age, and lifestyle factors, including diet, smoking and alcohol consumption. Approaches to quality control and imputation of the genotype data of UK Biobank have been described previously^7^, which filtered the samples to the European-ancestry only; similarly, the quality control and imputation of the genotype data of INTERVAL has been described in the previous work^4^. For algorithmic purposes, any remaining missing genotypes were mean imputed.

### Variant Selection and Interaction Detection

To construct PGSs for blood cell traits, a key step is to select genetic variants (e.g. SNPs), that are not only significantly associated with the trait but also independently contribute to the trait. Thus, a GWAS was first performed for each trait on the UKB cohort to select variants significantly associated with the trait, in which a MAF threshold of 0.005% was applied to ensure there are enough minor alleles per variant in the population for association tests; an INFO threshold of 0.4 was used to guarantee the confidence in genotype imputation accuracy, and a p-value threshold of 8.31×10^−9^ was used to identify significantly associated variants. Based on these significantly associated variants of each trait, a conditional analysis (CA) with a *r*^*2*^ threshold of 0.9 was further performed to identify the variants that are independently associated with a trait and can best represent the underlying genetic signals of that trait.

The conditional analysis was performed using a stepwise multiple linear regression approach^4,25^. For each blood cell trait, the set of genome wide significant variants was first partitioned into the largest number of blocks such that no pair of blocks are separated by fewer than 5Mb, and no block contains more than 2,500 variants. For each block, variants within the block are tested separately using the multiple-stepwise regression algorithm and independently associated variants are put forward into a larger chromosome wide pool on which a second multiple-stepwise regression algorithm is executed. The multiple-stepwise regression algorithm starts by adding in variants that pass the genome-wide significance threshold (8.31×10^−9^) and have a LD r^2^ score lower than 0.9. Then, it fits a multivariate linear regression to remove variants that have a p-values larger than the genome-wide significance threshold, which step is iterated until no more variants can be removed from the model. Note that we only keep those CA variants whose genotype data are available on both UKB and INTERVAL studies for the convenience of external tests in this study.

To investigate capability of learning methods in modelling interactions, variants interaction tests were performed on all the pairs of CA variants of a trait on the UKB cohort using multivariate linear regression: y = β_0_ + β_1_SNP_1_ + β_2_SNP_2_ + β_3_SNP_1_SNP_2_, and the interaction terms with p-value < 1×10^−8^ are deemed as valid variants interactions.

### Polygenic Scoring Methods

We constructed PGSs for 26 blood cell traits using a conventional univariate method as well as a variety of widely used machine learning and deep learning methods. This subsection describes fundamental aspects of the univariate (UNI), elastic net (EN), Bayesian ridge (BR) and multilayer precepton (MLP) and convolutional neural network (CNN) methods.

#### Univariate Method(UNI)

UNI method assumes that the genetic variants have linear additive effects on PGSs of the trait and constructs polygenic scores of a blood cell trait using the weighted sum of genotypes of the selected variants for that trait^13^:

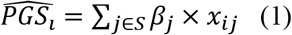

where *S* is the set of SNPs that are identified in the variants selection step; *β*_*j*_ *is* the effect size of the SNP *j* that is obtained through the univariate statistical association tests in the GWAS using the UKB cohort; *x*_*ij*_ is the genotype dosage of SNP *j* of the individual *i*.

#### Elastic Net(EN)

EN also assumes that the variants have linear additive effects on the PGS of a trait, i.e. Eq. (1), but the effect sizes of variants are obtained using a different way. These effect sizes are estimated by minimizing the penalized squared loss function:

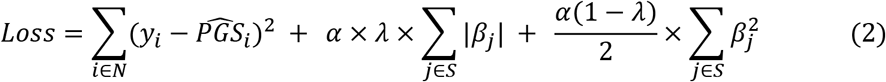

in which, *N* is the set of training samples for a given trait; the second term is L1 norm and the third term is L2 norm; α and λ are coefficients used to control the contribution of L1 and L2 norms in the model, which are usually set via cross-validation. In EN, effect sizes of the variants selected for a trait are jointly estimated which provides an implicit way to model the mutual interactions among these variants, and the use of L1 and L2 norms helps to control model complexity to address the over-fitting problem in which L1 controls the sparsity of the model and L2 controls the contribution of each variable. It has been shown that the application of these regularized multivariate models offers an effective way to improve PGS construction in practice^19,20^.

#### Bayesian Ridge(BR)

Similarly, BR also has a linear assumption for the effects of the variants, i.e. Eq.(1). Different from EN, BR assumes that PGSs of a trait follow a Gaussian distribution, and the prior for effect sizes of variants is also given by a spherical Gaussian:

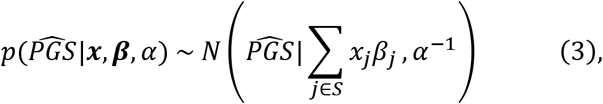

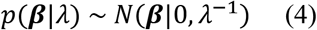

where α and λ are coefficients of the model and subject to two Gamma distribution: Gamma(α_1_, α_2_) and Gamma(λ_1_, λ_2_). These two prior Gamma distributions can be set via a validation step. The **β**, α, λ are then estimated by maximizing the log of the corresponding posterior distribution with respect to **β** by combining Eq. (3) and Eq.(4) on the training data^26^.

#### Multilayer Perceptron(MLP)

MLP is also named Deep Forward Neural Networks. Unlike other statistical learning methods, e.g. EN and BR, MLP makes no prior assumptions on the data distribution and can be trained to approximate virtually any smooth, measurable functions including non-linear functions^27^. A MLP typically consists of many different functions (or neurons) which are composed through a directed acyclic graph^28^. **Figure 2** shows an example of a three-layer MLP in which the first layer is known as input layer consisting of the input features, i.e. SNPs in the context of this study; the last layer outputs the final result of the model and the layer(s) in between are called hidden layer(s). A function node in hidden and output layers typically transforms the inputs from the previous layer with a weighted linear sum followed by an activation function^23^. For example, *f*^*1*^ in **Figure 2** can be represented as:

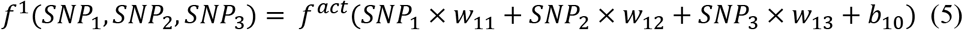

where *w*_*11*_, *w*_*12*_ and *w*_*13*_, are weights of the three inputs of function *f*^*1*^ and b_10_ is the intercept (or bias); SNP_1_, SNP_2_ and SNP_3_ are the genotype dosages of three SNPs in our context; *f*^*act*^ is an activation function which typically plays the role of introducing non-linearity into the model. Thus, the network architecture and its components of an MLP, e.g. activation function, determine a linear/non-linear mapping space, from which a model, i.e. all the weights across the given network that can best represent the data, is supposed to be learned. Details on the selection of network architectures for this study are given in the next subsection. This learning process is typically implemented by minimizing the difference, i.e. cost function, between the training data and the model distribution, through a back-propagation algorithm^28^.

**Figure 2.**
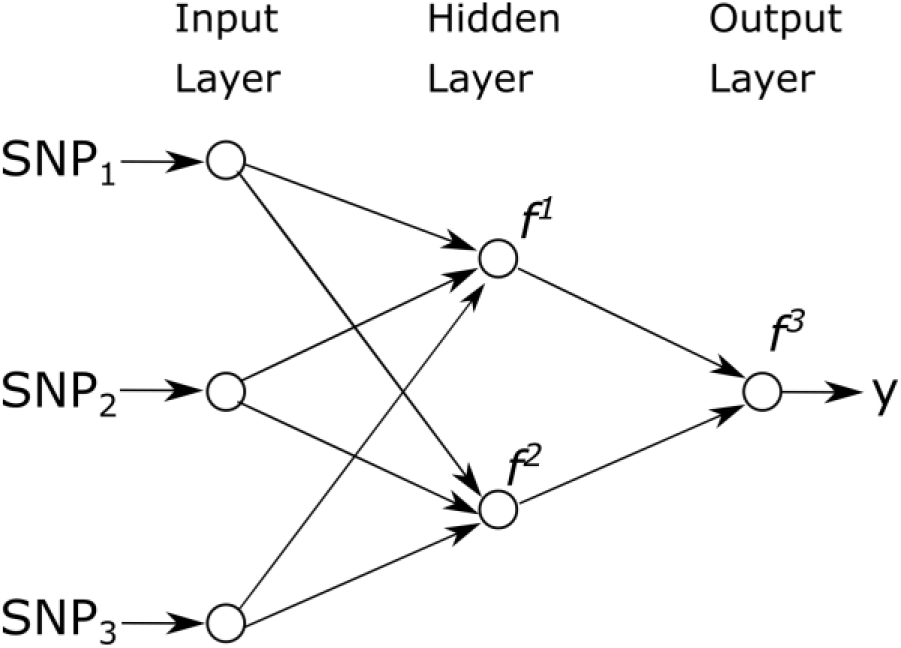
An example of a three-layer MLP. The output *y* = *f*^*3*^ (*f*^*1*^(*SNP*_*1*_, *SNP*_*2*_, *SNP*_*3*_), *f*^*2*^(*SNP*_*1*_, *SNP*_*2*_, *SNP*_*3*_)).

#### Convolutional Neural Networks(CNNs)

CNNs are a specialized neural network for processing data that have a grid-like topology^28^, e.g. time-series data, image data, genome sequence data^29^. As regularized versions of MLPs, CNNs construct its hidden layers using convolutional and pooling operations which are usually followed by fully connected layers and the output layer. The convolution operation limits the number of input units for an output unit by using kernels, and leads to a sparse connectivity of the network, which allows us to store fewer parameters and largely improve statistical efficiency. A typical convolutional layer in CNNs performs multiple convolutions in parallel which lead to multiple representations of the input units. To help generalise these representations and reduce the chance of overfitting, a pooling layer is usually followed to replace each representation at a certain location with a summary statistic of the nearby output units^28^. There are different pooling operations that can be applied based on different application context, e.g. max pooling and average pooling. **Figure 3** shows an example of a simple one-dimensional CNN with illustrations on convolution and pooling operations.

**Figure 3.**
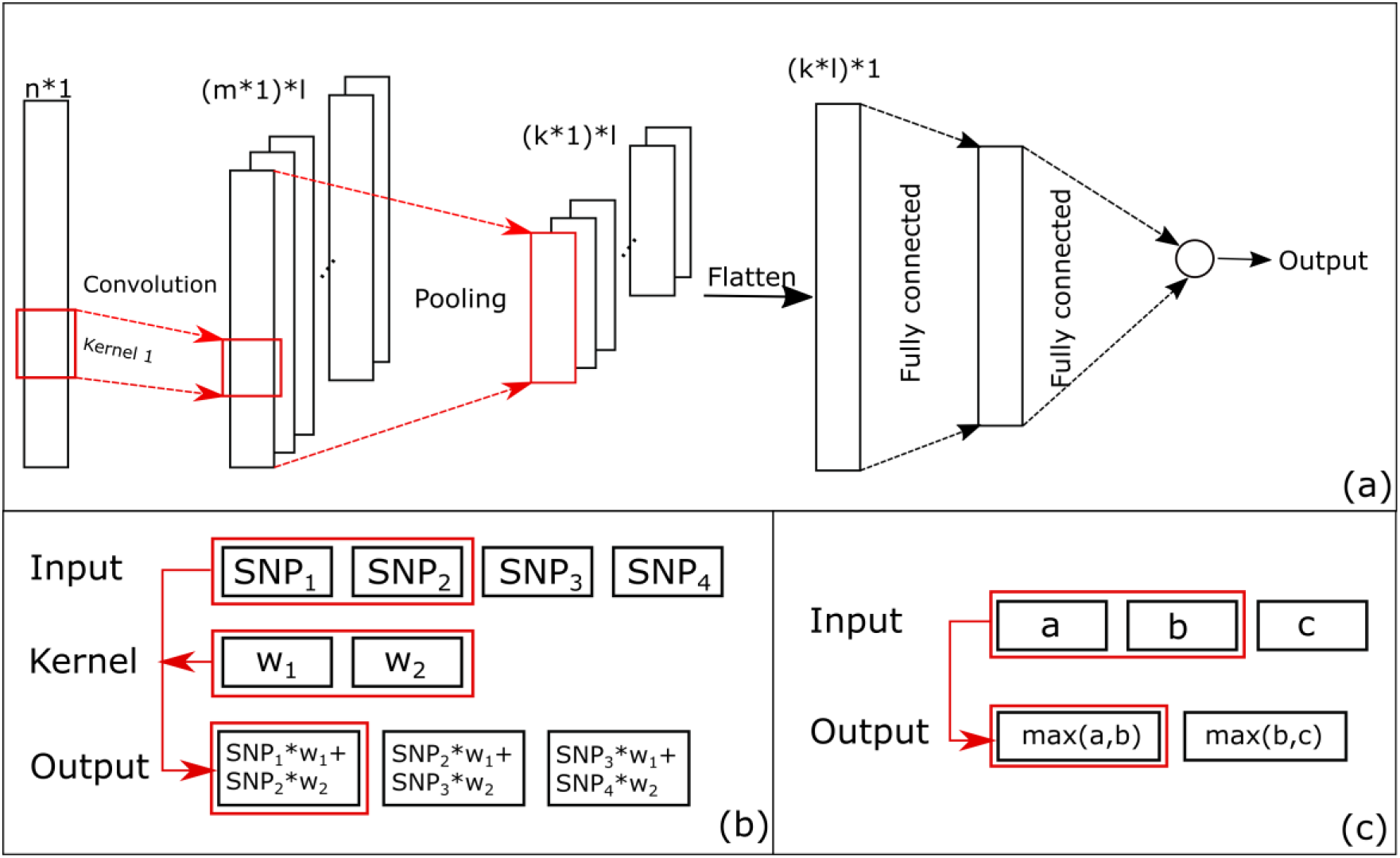
(a) An example of a one-dimensional CNN. (b) An example of a convolution operation. (c) An example of a max pooling operation. The convolution kernel in (b) has a size of 1*2 and operates with a stride of 1. The max pooling filter in (c) has a size 1*2 and operates with a stride of 1. The CNN in (a) has an input of a one-dimensional vector with *n* units, and has a convolution layer and a pooling layer. The dimension *m* of a newly generated representation via a convolution operation relies on the size of the kernel being applied as well as other possible factors, e.g. padding approaches, and the number of new representations *l* is equivalent to the number of kernels used in the model. The dimension *k* of a new representation after pooling is decided by the filter size being used.

### Measurement and Hyperparameter Tuning

We used Pearson ***r*** to measure the performance of various polygenic scoring methods. For each trait and each learning method, we randomly and equally partitioned the UKB samples into 5 portions, from which any 4 portions (80% of the samples) are used as training data to learn a model, and test the respective model’s performance on the remaining 20% of UKB samples, as well as an external validation using the whole INTERVAL cohort. For each learning method and each trait, we obtained 5 different models, each with a performance measurement for both the internal UKB test and the external INTERVAL test. By doing so, the training and internal testing covered the whole UKB cohort, affording an effective way to avoid evaluation bias. The UNI method was also tested on the five different UKB testing sets, and the whole INTERVAL cohort.

Hyperparameters turning is a crucial step for machine learning and deep learning methods as the choice of hyperparameters can greatly influence the model performance. In this study, a 10-fold cross-validation was performed on the training data to select the two hyperparameters α and λ of EN. To identify two appropriate gamma distributions in BR i.e. the selection of α_1_, α_2_, λ_1_ and λ_2_, a grid search across the set [−10^10^, −10^5^,−10, 0, 10, 10^5^, 10^10^] was conducted on the training set in which 10% of the samples were used as a validation set. EN and BR are implemented using the scikit-learn package (scikit-learn.org). As this work is, to our knowledge, the first attempt to employ MLPs and CNNs for genomic prediction of blood cell traits, there was no prior information that could be used for the design of network architecture for this task. Therefore, similar to the previous work^23^, we used a genetic algorithm to search for the optimal MLP and CNN architectures as well as other hyperparameters, e.g. the number of layers, the number of neurons at each layer, activation functions, optimizers, dropouts, etc., on the train set, in which 10% of the samples were used as a validation set. MLPs and CNNs were implemented using Keras (keras.io).

### Derivation of PGS for disease on INTERVAL

The polygenic risk score used for coronary artery disease (CAD) was our previously published CAD meta-GRS^30^; a polygenic score comprising 1.75 million variants derived from a meta-analysis of three PGSs for CAD in UK Biobank. Briefly, the three meta-analysed CAD PGSs were: (1) an earlier PGS^31^ comprising 46,000 metabochip variants and their log odds for CAD in the 2013 CARDIoGRAMplusC4D consortium GWAS meta-analysis^32^; (2) a PGS comprising 202 variants whose association with CAD in the 2015 CARDIoGRAMplusC4D consortium GWAS meta-analysis^33^ were significant at a false discovery rate (FDR) < 0.05; and (3) a genome-wide PGS derived from the same summary statistics^33^ LD-thinned at r^2^=0.9 threshold in UK Biobank (version 2 genotype data, imputed to the HRC panel only).

PGSs for schizophrenia, Crohn’s disease, rheumatoid arthritis, allergic disease and asthma were derived from summary statistics from their respective genome wide association studies (GWAS) by filtering to variants that overlapped with a set of 2.3 million linkage disequilibrium (LD)-thinned (r^2^ < 0.9), high-confidence (imputation INFO score > 0.4), common (MAF > 1%), unambiguous SNPs (A/T and G/C SNPs excluded) in the UK Biobank version 3 genotype data^6,34^ (imputed to the 1000 genomes, UK10K, and haplotype reference consortium (HRC) panels^35^). GWAS summary statistics used for schizophrenia, Crohn’s disease, rheumatoid arthritis, allergic disease, asthma, and lung function were those published in the previous works^36–40^.

Levels of each PGS in each INTERVAL participant were calculated using the UNI method implemented in plink version 2.00^41^. In the case of missing genotypes, the frequency of the effect allele in INTERVAL was used in its place. For each PGS, these total sums were subsequently standardised to have mean of 0 and standard deviation 1 across all INTERVAL participants. Variants with complementary alleles (*e.g.* A/T and G/C variants) were excluded to avoid incorrect effect allele matching due to strand ambiguity. Variants with INFO < 0.3 were removed. Where there were duplicate variants the one with the highest INFO score was kept. In total, 54,069,889 variants passed QC for PGS calculation of these diseases.

## Results

We first compared the performance of the four learning methods with that of the univariate method for constructing PGSs for 26 blood cell traits (**Figure 4**). The three learning methods (EN, BR and MLP) consistently outperform the univariate method in terms of Pearson ***r*** for nearly every blood cell trait. Notably, the performance of EN and BR were nearly indistinguishable and were the top performing methods overall. With any of the three learning methods, PGSs for 11 blood cell traits achieved a nearly 2% or more increase in Pearson *r* in internal tests. The following five blood cell traits each achieved about or more than 2% improvement in both internal and external tests using any of the three learning methods, in comparison with the UNI method: monocyte percentage (MONO%), white blood cell count (WBC#), mean platelet volume (MPV), monocyte count (MONO#) and plateletcrit (PCT). We found that incorporation of nonlinearity factors, as in MLP and CNN, did not improve genomic prediction of blood cell traits, compared with linear models. For nearly half of blood cell traits we studied, the CNN resulted in PGS with approximately the same or lower Pearson *r* as the univariate approach.

**Figure 4.**
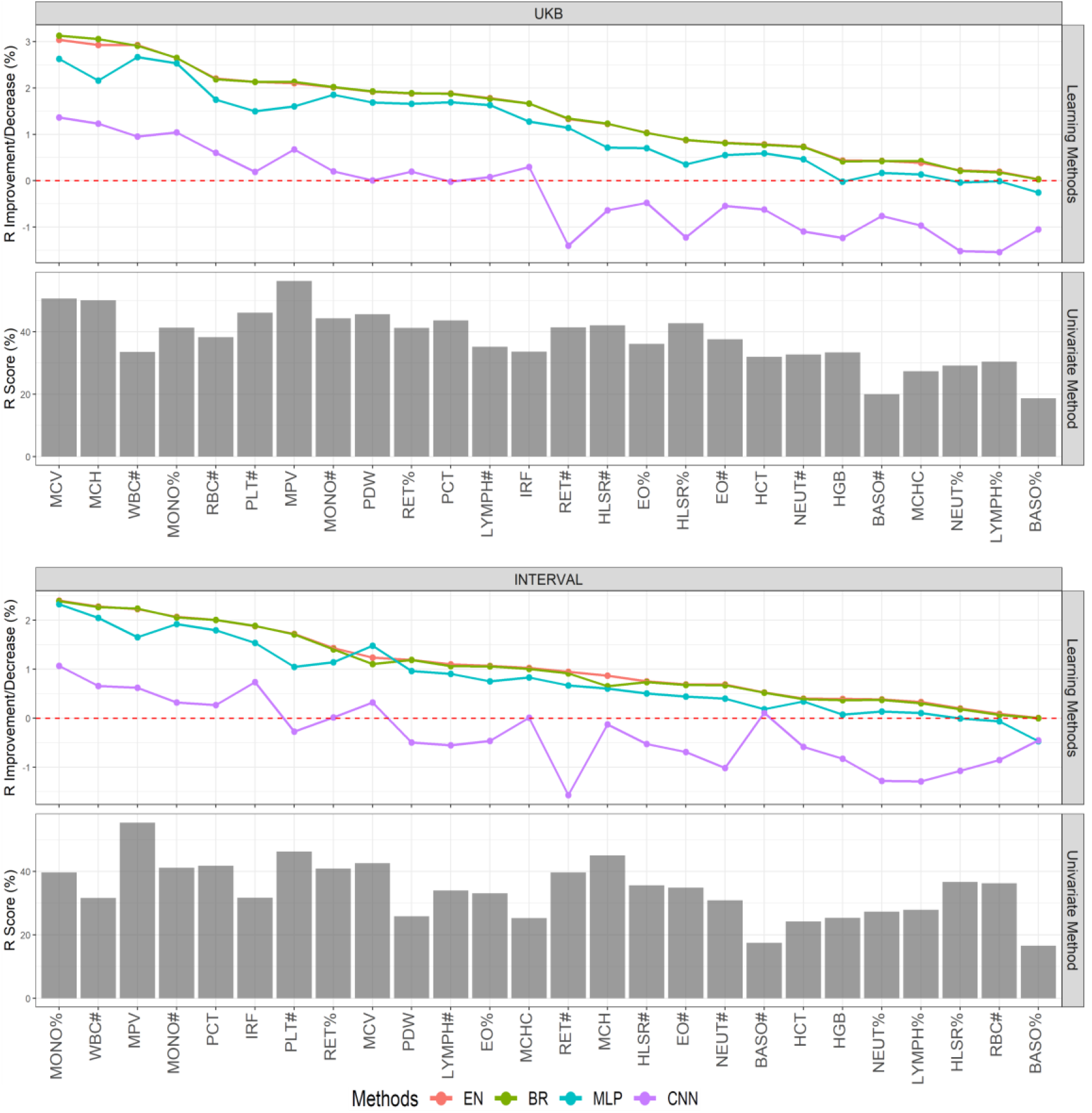
Performance comparison of different learning methods with the univariate method. Pearson *r* score performance of the UNI method for PGS construction of 26 blood cell traits are presented in testing on UKB or INTERVAL. Relative to the UNI method, performance of the four learning methods: EN, BR, MLP and CNN, are presented for each blood cell trait in descending order left to right according to EN (largest Pearson *r* increases on left). Given a particular method, a trait and a cohort, the averaged *r* performance of the 5 trained models, corresponding to the 5 different training-testing data partitions, is shown.

BR and EN outperformed UNI due largely to differences in variant effect size estimation **(Figure 5** and **Figure S1)**. We found that no effect sizes were set to zero by BR and EN, and effect sizes of most variants using BR or EN are the same or similar to those of the UNI method. This is consistent with a model where most common genetic variants are independently and additively contributing to each blood cell trait. Additionally, we also found both EN and BR tended to shrink the effects (sometimes greatly) of variants with low MAF; however, this did not necessarily contribute substantially to improved PGS. For example, effects of numerous low-MAF variants for traits like WBC# and hematocrit (HCT) were substantially shrunk by BR and EN; PGS construction of WBC# achieved significant improvement (~2% increases in Pearson *r*), while PGS for HCT saw little improvement. The effect shrinkage of low-MAF variants mainly resulted in better model generalization since many training samples did not have minor alleles of these low-MAF variants.

**Figure 5.**
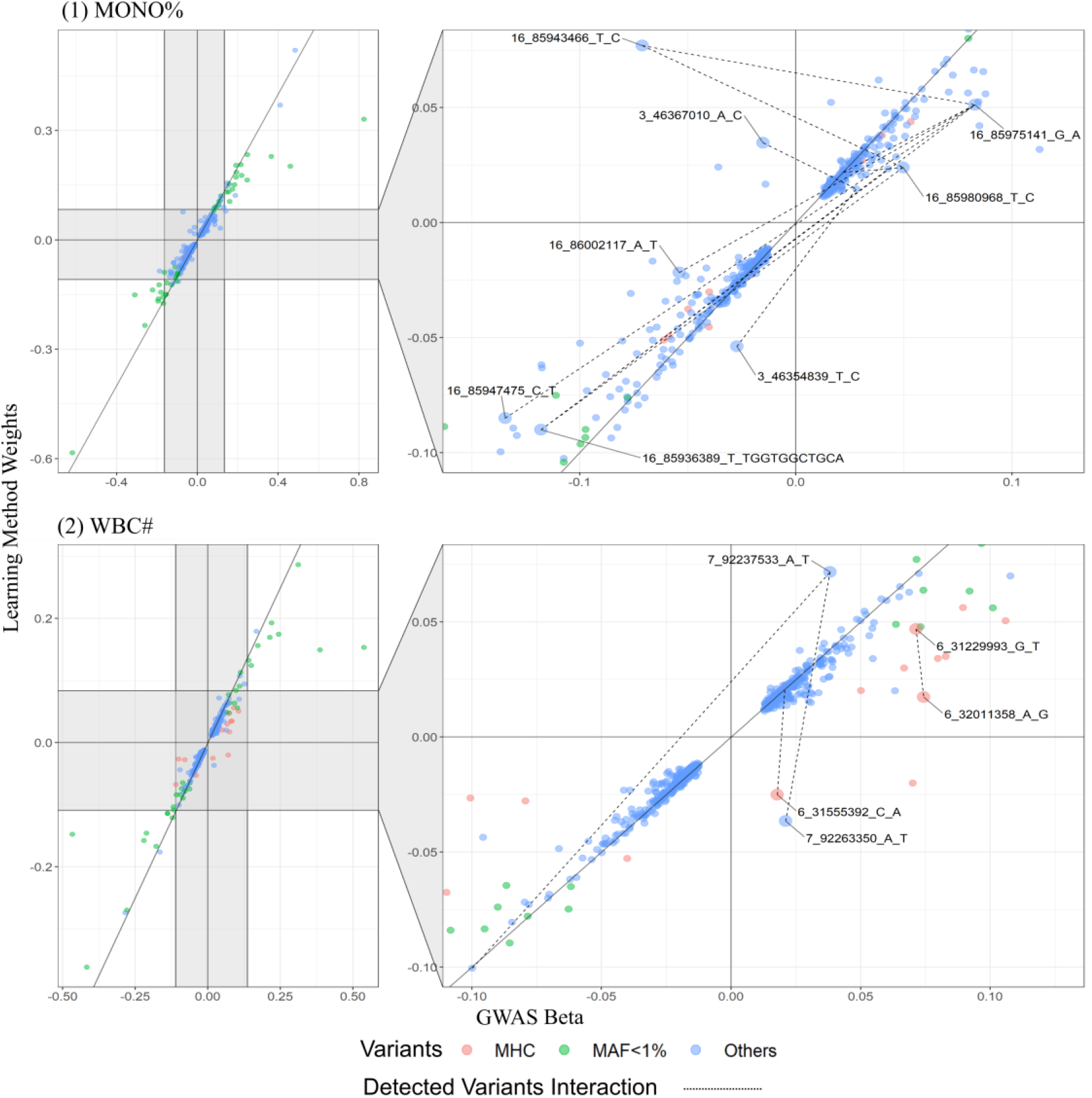
Comparison of variants effect sizes from UNI and BR method for trait MONO% and WBC#. EN and BR generated almost the same effect sizes for the variants of all the traits, thus for simplicity, this figure only compares the variants effect sizes between BR and UNI. The mean of the 5 effect sizes in the 5 trained BR models for each variant is used as the variant effect size of BR in this figure. Those variants from MHC region (25Mbp to 33Mbp on chromosome 6) are marked in red and the variants whose MAF is smaller than 1% are marked in green. Other variants are marked in blue. Those variants are both in the MHC region and have MAF<1% are considered as MAF<1% variants. Left section of the figure shows the overall distribution of all the variants of a blood cell trait. The right section is a zoomed-in area of the left section and its x-axis and y-axis only cover the ranges of these variants that have been detected with epistatic interactions. The two variants of each detected interaction of a trait are connected using dotted line on the right section of the figure. Those variants, whose effect sizes have the largest difference (top 10%) between the UNI and BR methods and are detected with epistatic interactions, are enlarged and marked with a variant identifier in the format of *chromosome number*, *base pair position*, *reference allele* and *alternative allele*.

We found that the superior performance of BR and EN was largely due to EN and BR changing the weights of variants with evidence of interaction effects on the blood cell trait **(Figure 5, Figure S1)**. For example, there were 12 significant interactions detected among 11 genetic variants closely located to each other on chromosomes 3 and 16 for trait MONO%, and effect sizes for most of these variants saw relatively large differences by EN and BR, with even some variant effects changing direction, compared to UNI. Similarly, there were 2 significant variants interactions detected in the MHC region and 2 interactions detected outside the MHC region for trait WBC#, and, as expected, effect sizes of these variants were substantially different in EN and BR compared to UNI.

To further demonstrate the role of interaction variants in PGS construction of blood cell traits, we removed the significant SNP-SNP interaction variants from the set of conditional analysis (CA) variants for each trait, and observed the *r* improvements using learning methods EN and BR, in comparison with UNI, on the pruned variants set on UKB and INTERVAL (Table S3). We found that the learning methods did not perform as well without the interaction variants compared with using the full CA variant set, for nearly every trait. For example, removing the 11 interaction variants for MONO% decreased the Pearson *r* improvement from 2.65% to 0.93% in UKB testing, and from 2.4% to 1.02% in INTERVAL; the removal of 7 interaction variants for WBC# reduced *r* improvement from 2.93% to 2.03% in UKB testing and from 2.28% to 1.5% in INTERVAL. These results demonstrated that the SNP-SNP interactions of a blood cell trait, among even a small set of variants, make significant contributions to its PGS, and the univariate PGS do not sufficiently capture these epistatic interactions^42^; whereas, data-driven learning may partially capture these epistatic effects by better adjusting combinations of marginal effects^20^.

Across nearly every blood cell trait we found that, even with prediction performance declining after removing interacting SNPs, BR and EN still saw consistent improvements compared with the UNI method. To test their robustness to redundant information, as would be expected from full genome-wide modelling, we allowed different sets of genetic variants to enter the model in addition to the variant set from conditional analysis (CA). We found that both EN and BR perform consistently well across the variants sets and all the blood cell traits, while the performance of UNI largely worsens after adding other variant sets (**Figure S2**). Overall, our results demonstrate the generally superior performance of learning methods, particularly BR and EN.

Maximising the accuracy of PGSs for blood cell traits raises opportunities for insights into underlying biology, potentially of relevance to disease risk. We next compared the extent to which EN-trained PGSs would be used to stratify the levels of blood cell traits in men and women over the age ranges of INTERVAL and UKB (**Figure 6** and **Figure S3**). There were a wide range of age-dependent dynamics in blood cell traits in both UKB and INTERVAL, with the EN-trained PGS offering stratification largely consistent with Pearson *r* of the trait. Blood cell traits exhibited well-known sex differences^43^. Interestingly, PGS for about half of blood cell traits showed different levels of stratification for men and women with 10 blood cell traits with a Bonferroni-adjusted significant interaction between sex and PGS (**Table 1**). For example, white blood cell indexes in women significantly decrease after menopause age, while the level of these traits in men were relatively stable^44^. Importantly, in both men and women, the EN-trained PGSs continued to stratify the trait levels even after the trait levels themselves changed. Average trait levels in the top versus bottom PGS quintiles were substantially different. The top quintile of the PGS for WBC# had an additional ~1.5 white blood cells per nanolitre (nL) on average in INTERVAL compared to the bottom quintile (an increase of ~25%); similarly, the difference between the top vs bottom 1% PGS for WBC# was ~2.2 white blood cells per nL (a 40% increase). For mean corpuscular volume, individuals in the top PGS quintile had red blood cells with ~5 femtolitres (fL) greater volume on average than those in the bottom PGS quintile, and these differences were maintained over all age ranges for both men and women.

**Figure 6.**
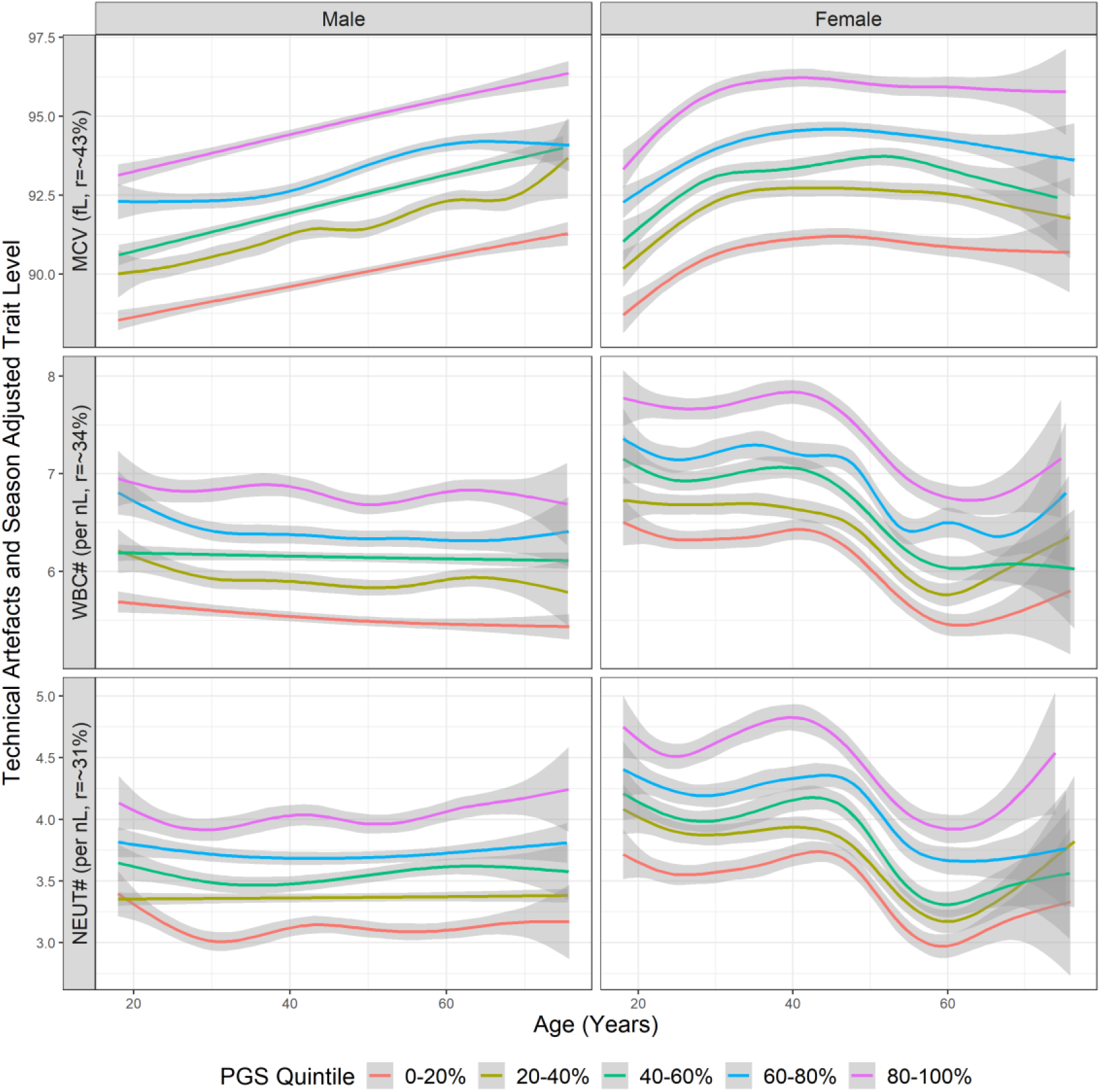
Trait levels by quintiles of EN-trained trait PGSs in men and women for trait MCV, WBC# and Neutrophil count (NEUT#) on INTERVAL. The shaded areas represent 95% confidence intervals.

**Table 1.**
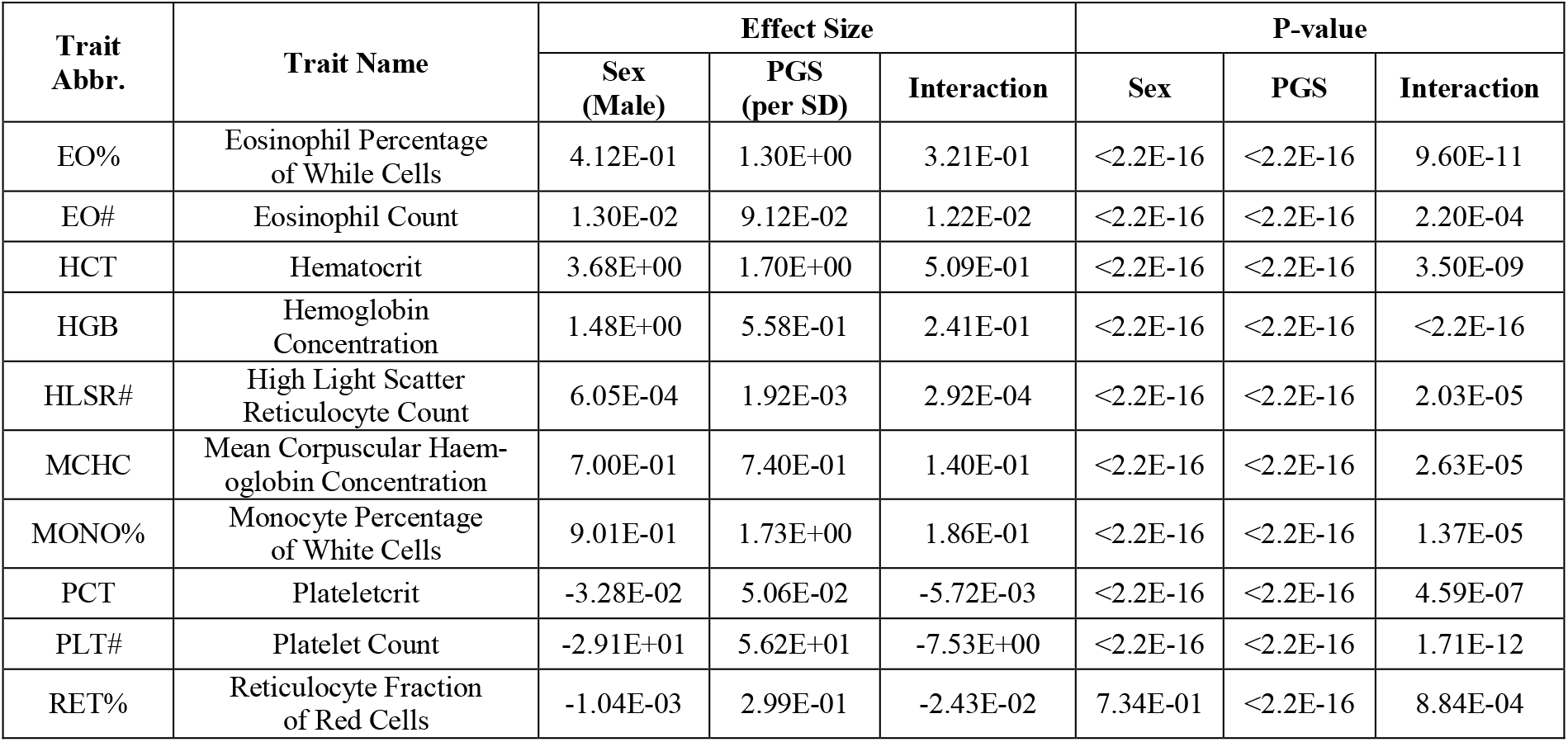
Summary statistics of PGS-Sex interaction tests for blood cell traits on INTERVAL. Interactions between PGSs and sex were tested for all the traits on the INTERVAL cohort by using the multivariate linear regression: y = β_0_ + β_1_*PGS + β_2_*Sex + β_3_*PGS*Sex, where y is the actual trait levels adjusted for technical artefacts, season, age and the first 10 genetic principal components; PGSs are construed using EN-trained models using UKB samples and standardised in the model. There are 10 traits whose p-values of interaction term passed the Bonferroni significance threshold 10^−3^, which are listed in the table. Standard deviation is abbreviated as SD.

| Trait Abbr. | Trait Name | Effect Size |  |  | P-value |  |  |
| --- | --- | --- | --- | --- | --- | --- | --- |
|  |  | Sex (Male) | PGS (per SD) | Interaction | Sex | PGS | Interaction |
| EO% | Eosinophil Percentage of White Cells | 4.12E-01 | 1.30E+00 | 3.21E-01 | <2.2E-16 | <2.2E-16 | 9.60E-11 |
| EO# | Eosinophil Count | 1.30E-02 | 9.12E-02 | 1.22E-02 | <2.2E-16 | <2.2E-16 | 2.20E-04 |
| HCT | Hematocrit | 3.68E+00 | 1.70E+00 | 5.09E-01 | <2.2E-16 | <2.2E-16 | 3.50E-09 |
| HGB | Hemoglobin Concentration | 1.48E+00 | 5.58E-01 | 2.41E-01 | <2.2E-16 | <2.2E-16 | <2.2E-16 |
| HLSR# | High Light Scatter Reticulocyte Count | 6.05E-04 | 1.92E-03 | 2.92E-04 | <2.2E-16 | <2.2E-16 | 2.03E-05 |
| MCHC | Mean Corpuscular Haemoglobin Concentration | 7.00E-01 | 7.40E-01 | 1.40E-01 | <2.2E-16 | <2.2E-16 | 2.63E-05 |
| MONO% | Monocyte Percentage of White Cells | 9.01E-01 | 1.73E+00 | 1.86E-01 | <2.2E-16 | <2.2E-16 | 1.37E-05 |
| PCT | Plateletcrit | -3.28E-02 | 5.06E-02 | -5.72E-03 | <2.2E-16 | <2.2E-16 | 4.59E-07 |
| PLT# | Platelet Count | -2.91E+01 | 5.62E+01 | -7.53E+00 | <2.2E-16 | <2.2E-16 | 1.71E-12 |
| RET% | Reticulocyte Fraction of Red Cells | -1.04E-03 | 2.99E-01 | -2.43E-02 | 7.34E-01 | <2.2E-16 | 8.84E-04 |

Finally, we examined the genetic correlations between the EN-trained PGSs of blood cell traits and PGSs of several common human diseases (**Figure 7**). We found many genetic correlations passing Bonferroni adjusted significance (p-value<10^−4^) and several were consistent with well-known associations between the blood cell traits themselves and the disease. For example, it is well known that asthma has a strong association with eosinophilic indices^4^, consistent with our analyses which show PGSs for EO# and EO% were strongly correlated with the asthma PGS. The strongest correlation of the schizophrenia PGS was with the PGS for WBC#, consistent with studies suggesting a potential correlation between WBC# itself and schizophrenia risk^45^. Our analyses also uncovered the shared genetic basis for previous trait-level observations for EO# and allergy disease^46^ as well as WBC# and Crohn’s disease^47^, while also demonstrating extensive shared polygenic basis for blood cell traits and rheumatoid arthritis, coronary artery disease, schizophrenia and Crohn’s disease, suggesting that, via shared genetics, blood cell traits may be either indicators or mediators for a variety of common disorders.

**Figure 7.**
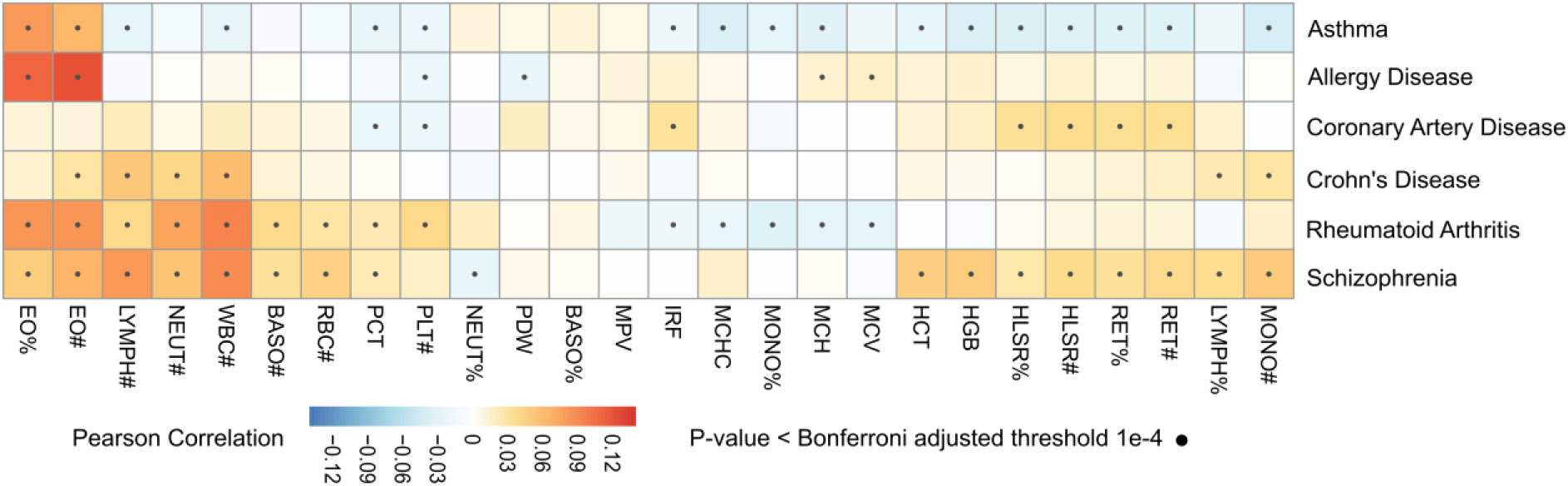
Correlation between PGSs for blood cell traits and PGSs for diseases in INTERVAL. PGSs for blood cell traits, diseases are adjusted for the first 10 genetic principal components before the calculation of correlations.

## Discussion

Improved polygenic models of blood cells traits promises to aid our understanding of myriad biological processes and diseases. This study demonstrated that machine and deep learning methods outperform conventional univariate methods in constructing polygenic scores for blood cell traits. In particular, our study’s results demonstrate that EN and BR methods can capture the effects of interactions among variants by adjusting their marginal effects in a linear model, thereby enabling these methods to outperform conventional approaches. This study also showed that PGSs are able to stratify age-dependent trait levels in both men and women in a population; that many blood cell trait PGSs have sex-specific interactions; and that there are extensive associations between PGSs of blood cell traits and various diseases. Collectively, these observations suggest the potential for these improved PGSs to help interrogate disease biology, enhance imputation of traits for association studies, and stratify populations into meaningful blood cell trait trajectories.

Our analysis indicates that EN and BR can partially capture interactions amongst genetic variants, resulting in improved PGSs. We observed consistent outperformance of EN and BR (compared with UNI) after removing the detected interaction variants, indicating the learning methods implicitly capture non-linear factors beyond these detected pair-wise interactions obtained using rigorous thresholds, e.g. multi-variants interactions and potentially interactions missed from univariately detected variants. The results also demonstrated the superiority of EN and BR in handling input noises, i.e. when incorporating redundant variants into the input variants set. This supports the use of loose input thresholds to incorporate more variants that may contribute to the trait, as the use of EN and BR would effectively dampen input noise allowed to enter the model.

Compared with EN and BR, MLP has a much looser model assumption, including the linear and non-linear models, but the increased model complexity did not result in improvements for PGSs construction of blood cell traits. In other words, the explicit incorporation of non-linearity factors in the model do not play a significant role for better PGSs construction of these trait as compared to EN and BR. This conclusion can also be supported by the results that the linear activation function was always selected in at least one of the top 10 MLP architectures (these models showed close performance on the validation data) for all the traits. On the other hand, it is also observed that many of activation functions in the selected top 10 MLPs is nonlinear for those traits showing significant improvements, e.g. MPV and WBC#, which hints at the existence of non-linear associations between the variants and the trait. Compared with MLP, CNN has a stricter model assumption and assumes an output unit in the network is only associated with the nearby input units. Its relatively poor performance (compared with other learning methods) in construction of PGS for blood cell traits suggests that this assumption could not fully capture the associations between the variants and these traits, and variants are associated with the trait in a more free manner. However, for both MLP and CNN, unlike the simple form of traditional machine learning models, the complex architecture of neural networks limits their interpretability.

This study demonstrated that populations can be well stratified by the PGSs of these blood cell traits, even for traits that exhibit quite large differences between ages and sexes. This shows the accuracy and robustness of the PGS construction method for these traits. By having accurate PGSs of these traits, it could help us to understand whether the existing differences in trait levels of an individual are due to genetic factors or/and other environmental factors in combination with sex, and furthermore, may allow us to identify individualized preventive or therapeutic targets. For example, it is known that some drugs, such as clozapine and dapsone^48^, have neutropenia side effects. The difference between top and bottom quintile of the neutrophil count PGS was ~1000 neutrophils per microlitre; therefore, there may be clinical utility in *a priori* knowledge that an individual may have genetically lowered neutrophil counts so as to guide pharmacotherapy.

The extensive sharing of the polygenic basis for blood cell traits and various common human diseases, was consistent with known trait-level associations and raised various new hypotheses. For example, both eosinophil count and neutrophil count are important risk factors for rheumatoid arthritis (RA), and their respective PGSs reflected these associations. Knowledge of their shared genetics and corresponding PGSs may enable early stratification of individuals at increased risk of eosinophil- or neutrophil-related RA. Such insights represent new avenues for using PGS to interrogate disease biology, and to facilitate that we have made the blood cell trait PGSs constructed here using EN available at the PGS Catalog^24^.

Overall, this study evaluated a variety of machine learning methods to construct PGSs for blood cell traits, highlighting the importance of moving beyond standard univariate methods and the capacity of learning methods to capture interactions amongst genetic variants. We make these PGS available to the community, and demonstrate that they can stratify sex- and age-dependent trajectories, and identify their shared polygenic basis with various common diseases. In the future, leveraging the totality of genetic variation for blood cell traits, as revealed in recent studies^49^, may represent further improvements of PGSs of these traits; Clinical uses of these PGSs will be another important focus of future studies.

## Acknowledgements

Participants in the INTERVAL randomised controlled trial were recruited with the active collaboration of NHS Blood and Transplant England (www.nhsbt.nhs.uk), which has supported field work and other elements of the trial. DNA extraction and genotyping was co-funded by the National Institute for Health Research (NIHR), the NIHR BioResource (http://bioresource.nihr.ac.uk) and the NIHR [Cambridge Biomedical Research Centre at the Cambridge University Hospitals NHS Foundation Trust] [*]. The academic coordinating centre for INTERVAL was supported by core funding from: NIHR Blood and Transplant Research Unit in Donor Health and Genomics (NIHR BTRU-2014−10024), UK Medical Research Council (MR/L003120/1), British Heart Foundation (SP/09/002; RG/13/13/30194; RG/18/13/33946) and the NIHR [Cambridge Biomedical Research Centre at the Cambridge University Hospitals NHS Foundation Trust] [*]. A complete list of the investigators and contributors to the INTERVAL trial is provided in reference [**]. The academic coordinating centre would like to thank blood donor centre staff and blood donors for participating in the INTERVAL trial.

This work was supported by Health Data Research UK, which is funded by the UK Medical Research Council, Engineering and Physical Sciences Research Council, Economic and Social Research Council, Department of Health and Social Care (England), Chief Scientist Office of the Scottish Government Health and Social Care Directorates, Health and Social Care Research and Development Division (Welsh Government), Public Health Agency (Northern Ireland), British Heart Foundation and Wellcome.

Dragana Vuckovic is funded by the National Institute for Health Research [Blood and Transplant Research Unit in Donor Health and Genomics] [*]. Scott Ritchie is funded by the National Institute for Health Research [Cambridge Biomedical Research Centre at the Cambridge University Hospitals NHS Foundation Trust] [*]. Tao Jiang was funded by the Cambridge University Medical Research Council Doctoral Training Programme. David Roberts is supported by NHS Blood and Transplant Research and Development funding and Oxford Biomedical Research Centre (Haematology Theme) [*]. John Danesh is funded by the National Institute for Health Research [Senior Investigator Award] [*] and British Heart Foundation [Personal Professorship].

## Supplementary Tables and Figures

**Table S1.**
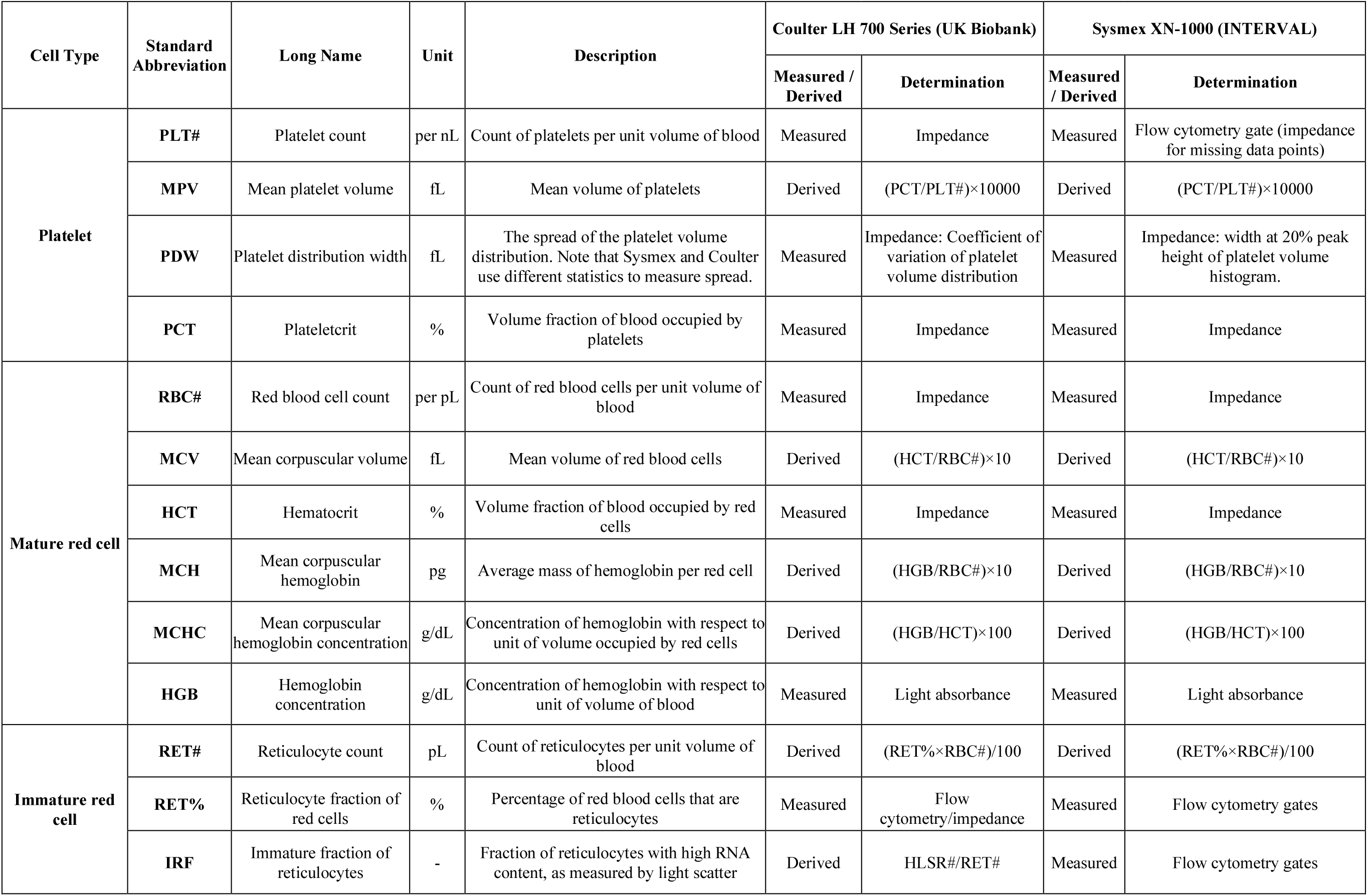
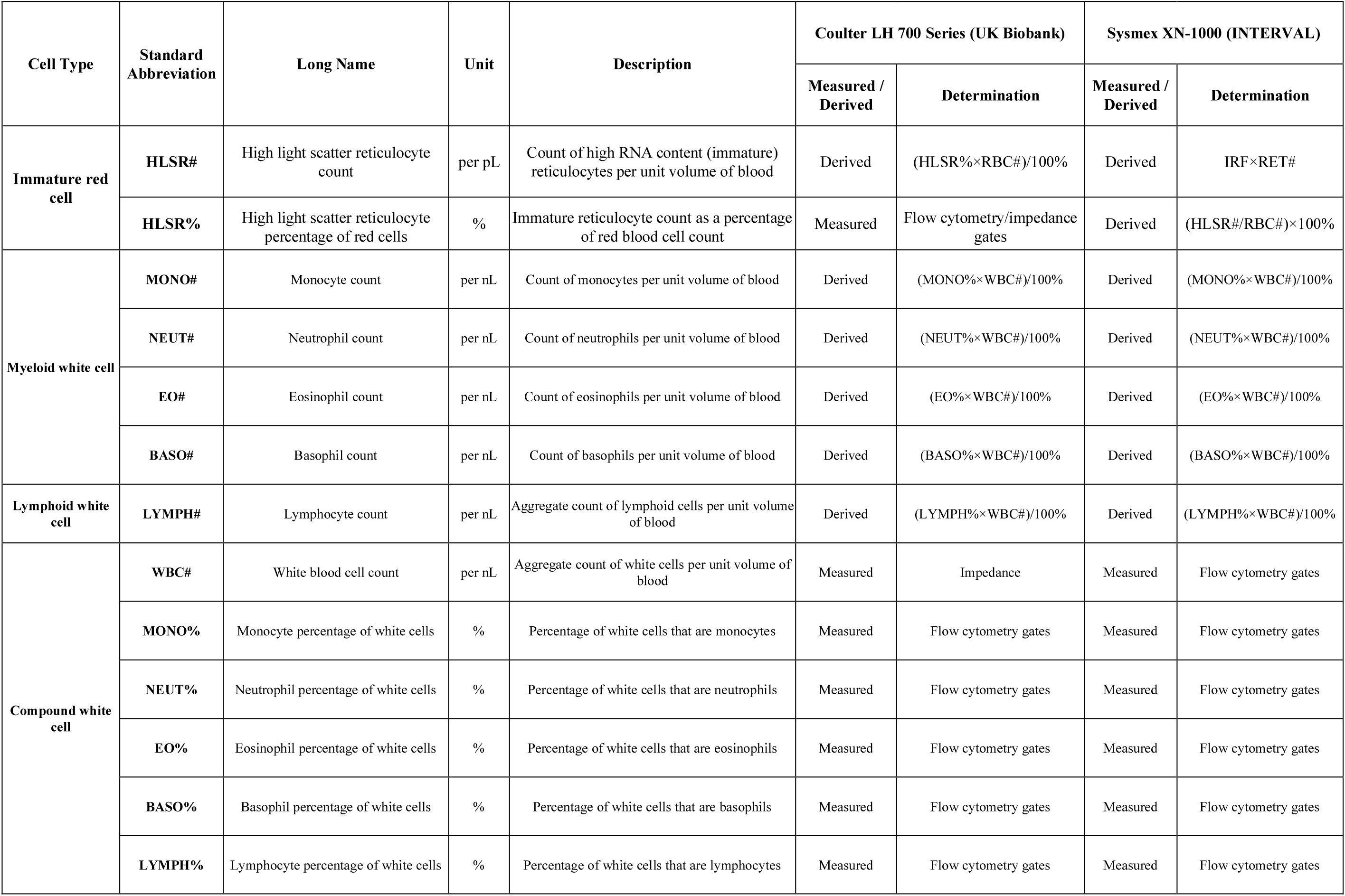
Summary of measurement methods for the 26 blood cell traits.

**Table S2.** The number of samples and variants used in UK biobank and INTERVAL for each blood cell trait. This table presents the number of valid samples for each trait after the quality control steps and the number of variants selected via conditional analysis.

| Trait | Number of Valid Samples |  | Number of Variants |
| --- | --- | --- | --- |
|  | UK Biobank | INTERVAL |  |
| PLT# | 391232 | 38939 | 762 |
| MPV | 391598 | 37224 | 681 |
| PDW | 391450 | 37262 | 579 |
| PCT | 390803 | 37306 | 726 |
| RBC# | 408069 | 40262 | 707 |
| MCV | 407157 | 40080 | 739 |
| HCT | 408112 | 40340 | 513 |
| MCH | 406517 | 40108 | 682 |
| MCHC | 407850 | 40265 | 252 |
| HGB | 407739 | 40329 | 532 |
| RET# | 396720 | 40253 | 590 |
| RET% | 396811 | 40286 | 572 |
| IRF | 396408 | 40227 | 390 |
| HLSR# | 400334 | 40244 | 605 |
| HLSR% | 400438 | 40225 | 594 |
| MONO# | 403994 | 39177 | 674 |
| NEUT# | 406788 | 39138 | 512 |
| EO# | 406470 | 40276 | 623 |
| BASO# | 404718 | 39986 | 198 |
| LYMPH# | 407277 | 39191 | 639 |
| WBC# | 408032 | 40466 | 659 |
| MONO% | 403136 | 39189 | 583 |
| NEUT% | 407114 | 39190 | 452 |
| EO% | 406417 | 40326 | 589 |
| BASO% | 404532 | 40133 | 160 |
| LYMPH% | 407319 | 39178 | 489 |

**Table S3.** R performance improvement comparison of PGSs construction for blood cell traits using learning methods with/without detected interaction variants in UK biobank and INTERVAL. The detected interaction variants were removed from the set of conditional analysis variants for each trait, and then the pruned variants set was used to evaluate the performance of UNI, EN and BR on UKB and INTERVAL (the same procedure described in the **Data and Methods** section). The table presents the gained *r* improvements using EN and BR, compared with UNI, on the full conditional variants set and the pruned variants set in UKB and INTERVAL.

| Traits | Number of Interactions | Number of Interaction Variants | UK Biobank |  |  | INTERVAL |  |  |
| --- | --- | --- | --- | --- | --- | --- | --- | --- |
|  |  |  | With Interaction Variants | Without Interaction Variants | Difference | With Interaction Variants | Without Interaction Variants | Difference |
| MONO% | 12 | 11 | 2.65% | 0.93% | 1.71% | 2.40% | 1.02% | 1.39% |
| MONO# | 13 | 15 | 2.02% | 0.79% | 1.23% | 2.07% | 0.92% | 1.15% |
| MCHC | 8 | 14 | 0.39% | 0.21% | 0.18% | 1.03% | 0.17% | 0.86% |
| WBC# | 4 | 7 | 2.93% | 2.03% | 0.90% | 2.28% | 1.50% | 0.78% |
| LYMPH# | 18 | 12 | 1.78% | 0.25% | 1.54% | 1.10% | 0.36% | 0.73% |
| MPV | 15 | 20 | 2.11% | 1.36% | 0.74% | 2.23% | 1.57% | 0.66% |
| PLT# | 6 | 11 | 2.13% | 1.70% | 0.43% | 1.72% | 1.21% | 0.51% |
| HCT | 3 | 5 | 0.79% | 0.20% | 0.59% | 0.40% | 0.13% | 0.27% |
| EO% | 4 | 8 | 1.04% | 0.75% | 0.29% | 1.07% | 0.84% | 0.23% |
| NEUT# | 3 | 5 | 0.74% | 0.50% | 0.24% | 0.69% | 0.49% | 0.20% |
| RET% | 1 | 2 | 1.88% | 1.82% | 0.06% | 1.43% | 1.25% | 0.18% |
| PCT | 5 | 8 | 1.88% | 1.61% | 0.27% | 2.00% | 1.84% | 0.17% |
| EO# | 5 | 9 | 0.82% | 0.56% | 0.26% | 0.69% | 0.52% | 0.17% |
| IRF | 4 | 5 | 1.66% | 1.11% | 0.56% | 1.88% | 1.74% | 0.14% |
| HGB | 1 | 2 | 0.43% | 0.39% | 0.05% | 0.39% | 0.26% | 0.13% |
| RET# | 4 | 7 | 1.33% | 1.05% | 0.29% | 0.95% | 0.89% | 0.06% |
| NEUT% | 1 | 2 | 0.23% | 0.11% | 0.11% | 0.38% | 0.36% | 0.02% |
| HLSR# | 1 | 2 | 1.22% | 1.22% | 0.00% | 0.75% | 0.74% | 0.01% |
| HLSR% | 1 | 2 | 0.88% | 0.87% | 0.00% | 0.20% | 0.19% | 0.01% |
| BASO# | 0 | 0 | 0.43% | 0.43% | 0.00% | 0.53% | 0.53% | 0.00% |
| BASO% | 0 | 0 | 0.03% | 0.03% | 0.00% | 0.00% | 0.00% | 0.00% |
| LYMPH% | 0 | 0 | 0.19% | 0.19% | 0.00% | 0.33% | 0.33% | 0.00% |
| PDW | 4 | 7 | 1.92% | 0.94% | 0.98% | 1.19% | 1.26% | -0.08% |
| MCV | 28 | 31 | 3.04% | 1.29% | 1.75% | 1.24% | 1.37% | -0.13% |
| MCH | 27 | 31 | 2.93% | 1.31% | 1.62% | 0.87% | 1.09% | -0.22% |
| RBC# | 6 | 6 | 2.21% | 0.81% | 1.39% | 0.09% | 0.83% | -0.74% |

**Figure S1.**
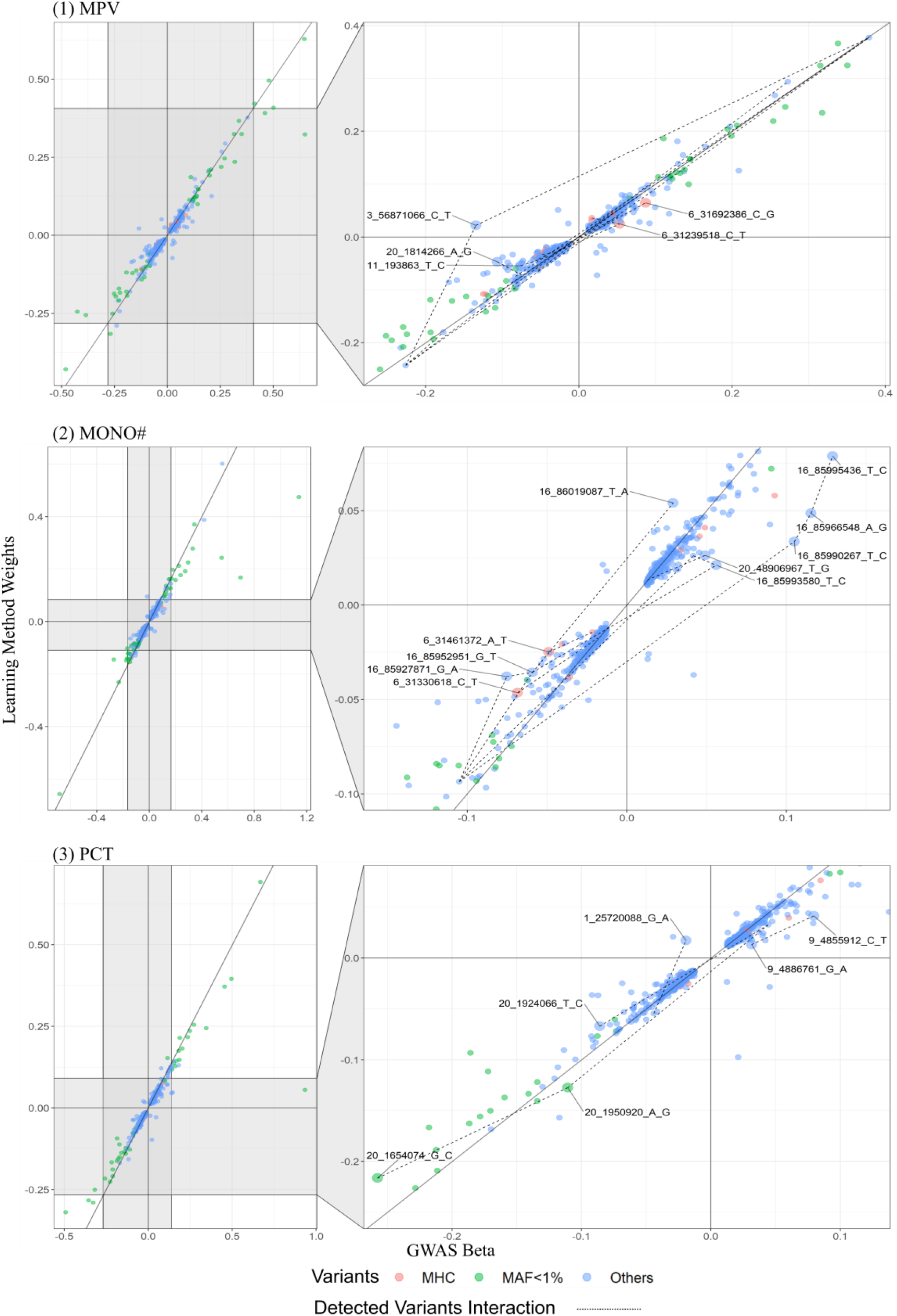
Comparison of variants effect sizes using UNI and BR methods for trait MPV, MONO# and PCT.

**Figure S2.**
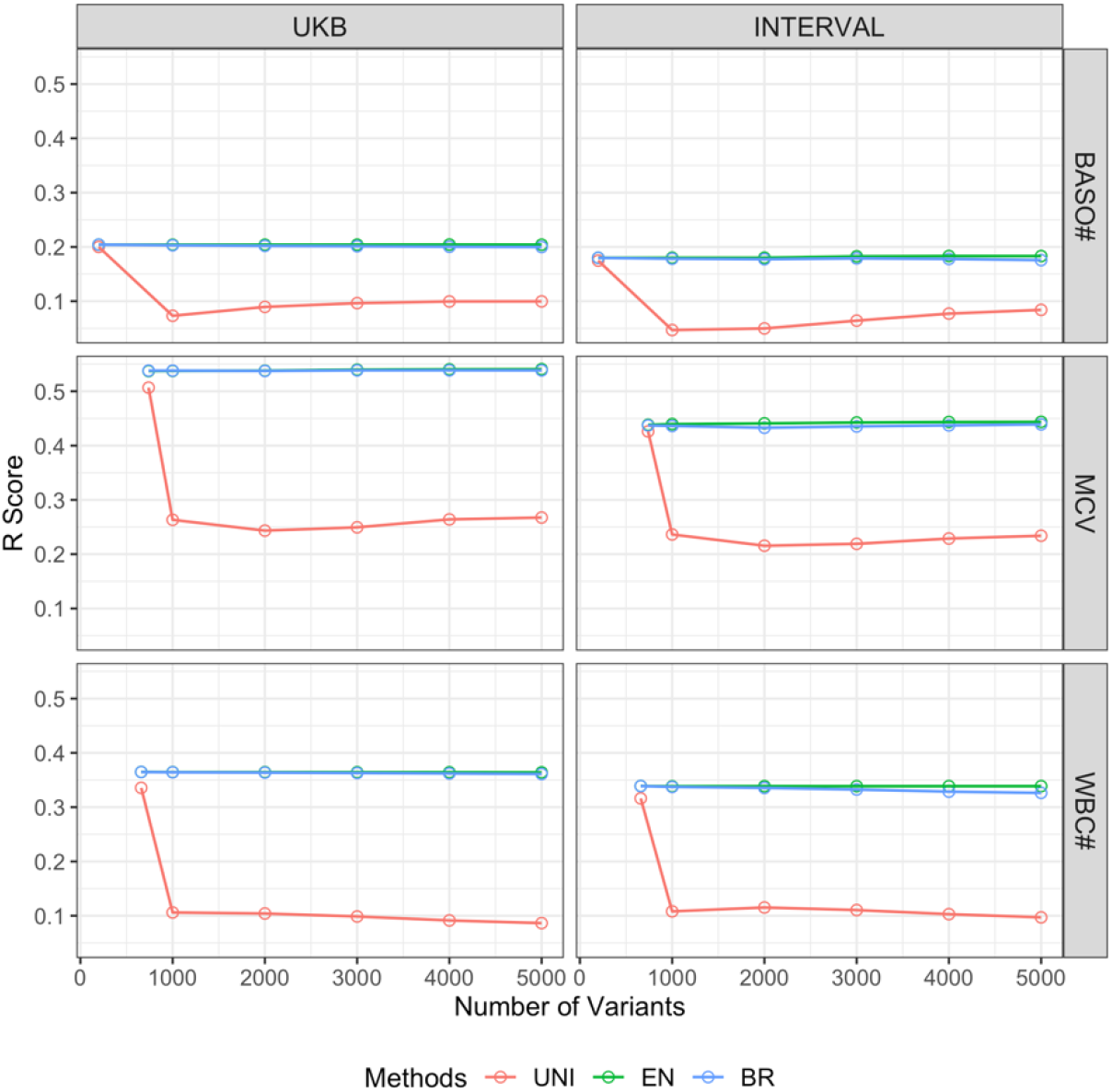
Performance of UNI, EN and BR methods on different sets of variants. Using CA variants as a base set, we increasingly add in the top significant variants by the step of 1000 (based on the GWASs on the UKB cohort) to form different sizes of variants sets, in which the CA variants are removed from the top significant variants list. It means the starting point of each curve is the *r* performance of the corresponding method on the CA variants set of a trait. We then observe the performance of BR, EN and UNI on different variants sets. Similarly, five models were trained corresponding to the five partitions of the UKB samples for each learning method, each trait and each variants set, whose *r* measurements are averaged in the figure. The results indicate that BR and EN are robust to redundancy in variants which appear to be through two different avenues: effect sizes of the CA variants generated by the two methods are very similar; however, EN tends to make the effect sizes of those non-contributing variants as zeros due to the use of L1 norm, while BR tends to make their effect sizes very small or can cancel each other due to the use of the L2 norm.

**Figure S3.**
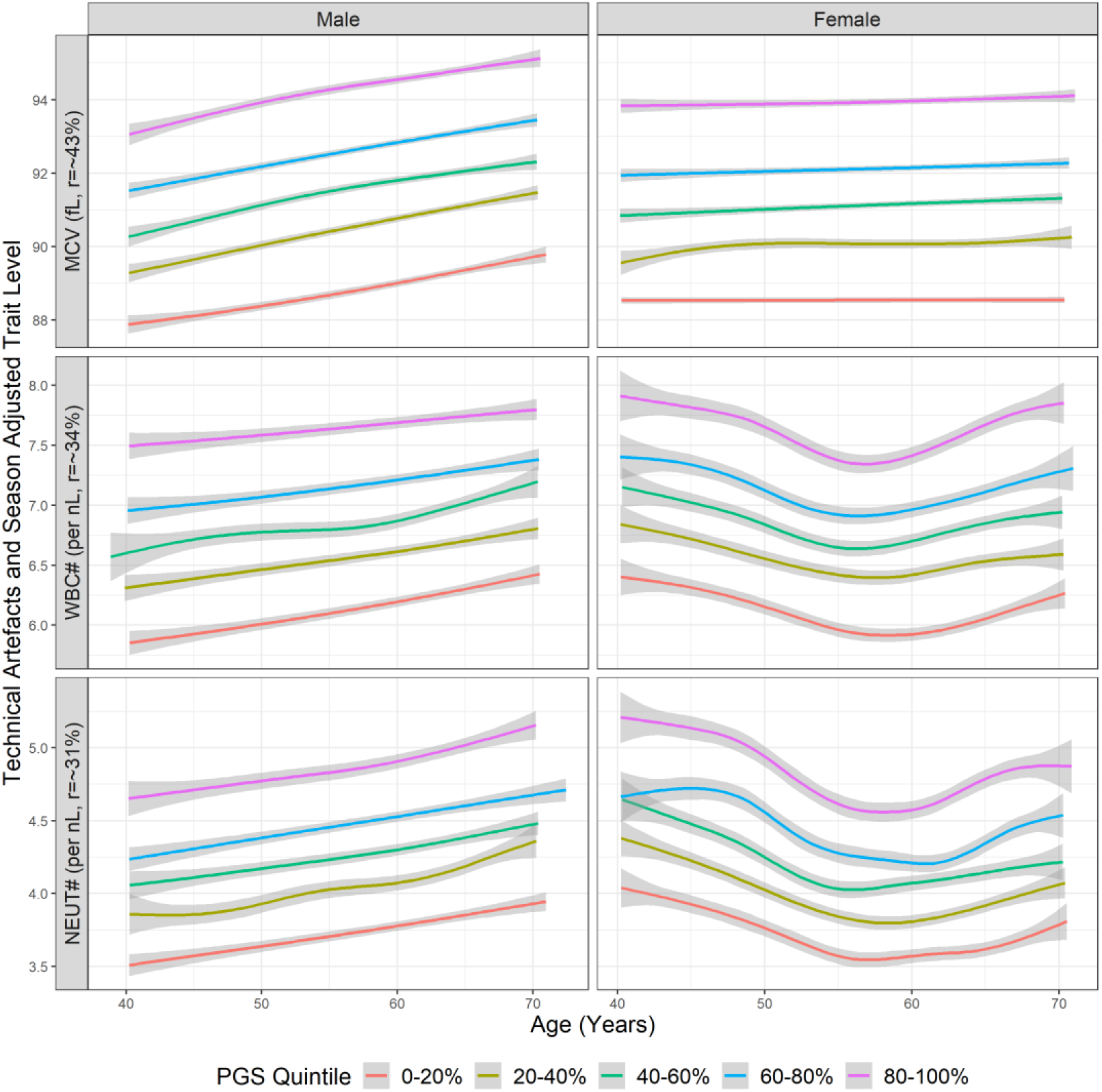
Trait levels by quintiles of EN-trained trait PGSs in men and women for trait MCV, WBC# and NEUT# on UKB.

## Footnotes

* The views expressed are those of the authors and not necessarily those of the NHS, the NIHR or the Department of Health and Social Care.

** Di Angelantonio E, Thompson SG, Kaptoge SK, Moore C, Walker M, Armitage J, Ouwehand WH, Roberts DJ, Danesh J, INTERVAL Trial Group. Efficiency and safety of varying the frequency of whole blood donation (INTERVAL): a randomised trial of 45 000 donors. Lancet. 2017 Nov 25;390(10110):2360-2371.

